# Darwin’s Naturalization Conundrum explained by gradients of environmental stress and disturbance

**DOI:** 10.1101/2024.10.17.618817

**Authors:** Andrea Galmán, Jake M. Alexander, Lohengrin A. Cavieres, Curtis C. Daehler, Tim Seipel, Marten Winter, Kelly M Andersen, Jonas J. Lembrechts, Valeria Aschero, José Ramón Arévalo Sierra, Agustina Barros, Meike Buhaly, Amanda Ratier Backes, Pervaiz A. Dar, Eduardo Fuentes-Lillo, Alejandra Jiménez, Christoph Küffer, Christian D. Larson, Maritza Mihoc, Ann Milbau, John W. Morgan, Keith McDougall, Jana Müllerová, Bridgett Naylor, Martin A. Nuñez, Miguel Antonio Padrón Mederos, Aníbal Pauchard, Lisa J. Rew, Zafar A. Reshi, Veronica Sandoya, Manzoor A Shah, James M. Shannon, Neville Walsh, Graciela Valencia, Tom Vorstenbosch, Genevieve Wright, Shengwei Zong, Sylvia Haider

## Abstract

Darwin’s Naturalization Conundrum (DNC) states that non-native species closely related to the native community are either more likely to succeed because shared adaptations help them overcome environmental filtering, or less likely to succeed because of strong competition with their native relatives. Despite extensive research, no general patterns have so far emerged. One reason may be that the relative importance of competition and environmental filtering depends on environmental conditions. To test this hypothesis, we conducted a global assessment of DNC examining patterns of phylogenetic relatedness of non-native plant species to the native community along gradients of elevation and anthropogenic disturbance in mountains. Phylogenetic distance of non-native to native species decreased with increasing elevation and in disturbed plant communities. Our results help resolve DNC by showing that the environmental context sets expectations for patterns of relatedness between non-native and native species and helps illuminate the ecological and evolutionary processes generating these patterns.

## INTRODUCTION

Plant invasions are one of the main drivers of global change, threatening biodiversity (Bellard et al. 2022, IPBES 2023), altering ecosystem functioning and leading to homogenization at different levels of biodiversity (Vilà et al. 2011, Winter et al. 2009). Understanding the ecological mechanisms that enable non-native species (i.e., species introduced beyond their native biogeographic range as a direct or indirect result of human action; Richardson et al. 2000) to succeed in novel environments is a central goal in ecology. Darwin (1859) was the first to propose evolutionary relatedness between non-native and native species as an important factor explaining invasion success and failure. This idea assumes that taxonomic or evolutionary (also referred to as phylogenetic) relatedness is a proxy for ecological similarity (Faith 1992, Webb 2000), and led to contrasting predictions about the role of relatedness for invasion success; this has been termed “Darwin’s Naturalization Conundrum” (DNC; Daehler 2001). Specifically, non-native taxa closely related to the native species in the novel environment are either (i) more likely to succeed because they share adaptations to the environment that help them overcome environmental filtering (pre-adaptation hypothesis; Ricotta et al. 2009), or (ii) less likely to establish due to strong competition with their native relatives (limiting similarity hypothesis or Darwin’s naturalization hypothesis; Torrey and Gray 1841, Rejmánek 1996). Several studies have addressed DNC, but results have been mixed, with positive, negative or no association between relatedness and establishment probability (e.g. Procheş et al. 2008, Thuiller et al. 2010, Jones et al. 2013, Maitner et al. 2021).

An important source of variability in studies addressing DNC may be that biotic competition and environmental filtering are not mutually exclusive (Cavender-Bares et al. 2006, Procheş et al. 2006), but can operate simultaneously or alternately depending on the context. For instance, the relative strength of these mechanisms has been found to depend on the spatial scale considered (Diez et al. 2008, Ma et al. 2016, Cadotte et al. 2018). Non-native species closely related to the native community were found to be more successful at regional scales, where environmental filtering is putatively of high importance (Swenson 2007), while closely related species experienced exclusion at local scales, where competitive interactions are more important (Procheş et al. 2008, Jiang et al. 2010, Peay et al. 2012). Just as the relative importance of environmental filtering and biotic interactions may vary with spatial scale, theory suggests that they may also vary with different environmental conditions (Kraft et al., 2015). Accordingly, a recent study showed that the presence of native species closely related to potentially invasive species is more likely to predict invasion success in harsher climates in the United States (Park et al. 2020). The influence of different environments on the strength and direction of environmental filtering and biotic interactions has been rarely evaluated in the context of DNC. Here we specifically explore how the effect of phylogenetic relatedness on invasion success varies with environmental conditions by exploring patterns of phylogenetic distance of non-native species to native species assemblages along two types of environmental gradients: elevational gradients and orthogonal gradients of anthropogenic disturbance.

Elevational gradients represent gradients of environmental stress, particularly climatic stress. The stress-dominance hypothesis (terminology adapted from Swenson & Enquist 2007) posits that competition is strong and provides a significant biotic filter for community membership under favorable environmental conditions, as typically found in more productive environments (e.g., low elevations in temperate mountain regions). Conversely, according to this hypothesis, the role of competition relative to abiotic conditions in shaping community membership is expected to be less important under environmental conditions that are more limiting to plant growth, such as low-productivity, high-elevation environments. Hence, while the relative role of competition is expected to decrease with elevation, the effect of environmental filtering is expected to increase, as fewer species with specific functional adaptations are able to thrive under such harsh conditions (e.g. Körner et al. 2003, Ratier Backes et al. 2023). Assuming that closely related species are functionally more similar (Webb 2000, Wiens et al. 2010, Swenson 2013), environmental filtering should generate communities with patterns of phylogenetic clustering (i.e., higher levels of relatedness among species than expected by chance, e.g., Prinzing et al. 2014). We can expect that non-native species will be more likely to invade environmentally stressful high elevation communities, if they are closely related to the resident native species. In contrast, in less stressful and more productive environments, characteristic of most low-elevations locations, competition could lead to patterns of phylogenetic over- dispersion, assuming that closely related and potentially functionally similar species competitively exclude one another. Simultaneously, the limited ecological niche overlap of distantly related species would reduce competition and facilitate species coexistence. However, there is increasing evidence that competition may also lead to phylogenetic clustering, if certain phylogenetic lineages are associated with competitively dominant traits (Mayfield & Levine 2010, Kraft et al. 2015).

The effect of anthropogenic disturbance represents another stressor that influences community assembly processes and the establishment of non-native plants (Lembrechts et al. 2016; Xu et al. 2020; here defined as the removal of plant biomass). Disturbed environments are characterized by setbacks of succession to earlier stages (Güsewell and Klötzli 2012), typically reducing competition for light, space or nutrients. Therefore, disturbance should allow phylogenetically similar non-native species to establish and co-exist with the native species. In general, non-native plant species have been found to benefit from reduced competition in disturbed habitats (e.g., Catford et al. 2012), especially under benign environmental conditions (Pauchard et al. 2009; Lembrechts et al. 2014). Elevation gradients along roads offer a special case for studying the interacting effects of environmental severity and disturbance on the establishment of non-native species, providing an opportunity to unravel the ecological processes driving non-native plant establishment. When comparing disturbed roadside environments and less disturbed natural environments adjacent to the road along elevation gradients, it can be expected that changes in the strength of competition and environmental filtering with increasing elevation will be more pronounced in natural environments than in disturbed roadside environments. If so, we hypothesize to find with increasing elevation a greater decrease in phylogenetic distance between non-native species and the native species community in natural environments, supporting the role of competition as a force promoting limiting similarity at low elevations.

In this study, we conducted a global assessment of Darwin’s Naturalization Conundrum (DNC) using vegetation survey data collected by the Mountain Invasion Research Network (MIREN; www.mountaininvasions.org; Haider et al. 2022) in 16 globally distributed mountain regions. We examined variability in phylogenetic distance between non-native and native species along environmental gradients as an approach to disentangle the relative role and importance of environmental filtering and biotic competition as processes determining non-native species establishment. First, we studied variability in phylogenetic distance patterns between non-native and native species within a plant community using vegetation surveys along elevational gradients as an approach to analyze the effect of environmental stress filtering non-native species. Second, we investigated phylogenetic distance patterns between non-native and native species by comparing plant communities adjacent to mountain roads (i.e., disturbed plant communities) and paired plots away from roads (i.e., more natural, less disturbed communities) to analyze the effect of disturbance and its interaction with elevation as a proxy for a reduction in competition. We use molecular-based phylogenies which provide a way to easily assess species’ relatedness in the context of genetic distance over time (Hughes and Eastwood 2006) in a more precise way than taxonomic approaches that use the number or abundance of congeneric species (Thuiller 2010). We hypothesized that (1) phylogenetic distance between non-native species and the native community should decrease with increasing elevation, assuming that environmental filtering selects for a phylogenetic subset of the non-native species pool, and that (2) phylogenetic distance between non-native and native species would be smaller in disturbed roadside habitats compared to less disturbed non-road habitats, assuming a weaker role of limiting similarity (competition) and a correspondingly stronger role of environmental filtering (Figure 1). Further, we expected that (3) phylogenetic distance between non-native species and the native community decreases more rapidly with elevation in undisturbed compared to disturbed communities (Figure 1d), assuming that reduced competition with increasing elevation and in disturbed environments leads to a reduction of limiting similarity.

**Figure 1:**
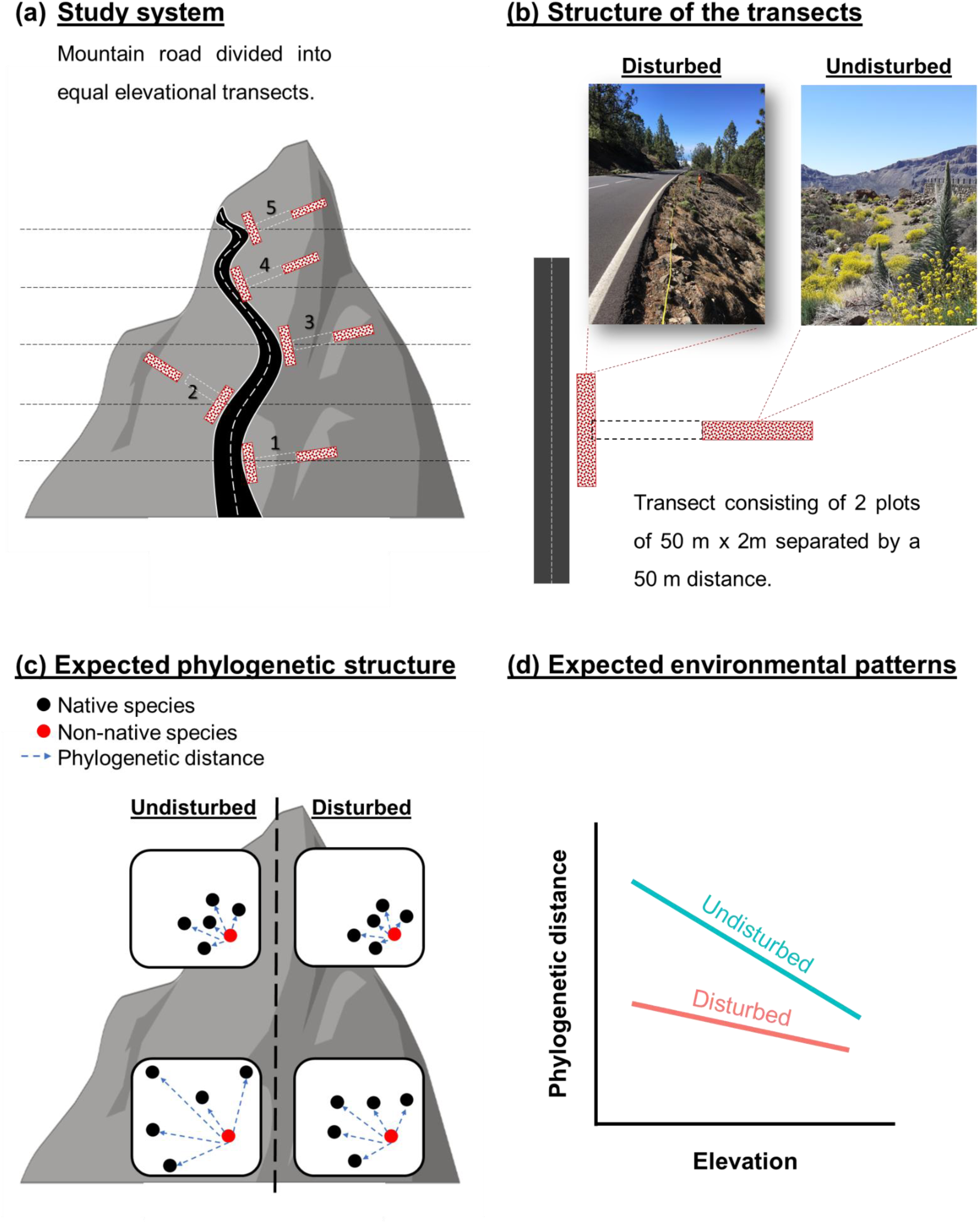
Conceptual figure of the study design and our hypotheses and predictions. Panel (a) shows an example of a mountain road with evenly spread sample sites (only five sample sites are shown for clarity). Panel (b) shows the structure of each sample site consisting of two plots of 50 m x 2m each, separated by a 50 m distance. The roadside plot (here considered as disturbed plot) is parallel to the road, the natural plot adjacent to the road (here considered as undisturbed) is centered 75 m from the roadside plot. Panels (c) and (d) show the hypotheses of the study: because of increasing environmental filtering and reduced competition towards harsher environments and in more disturbed habitats, we expect to find more clustered native species and a smaller mean phylogenetic distance of the non-native to the native species at high compared to low elevations and in disturbed compared to undisturbed communities. The differences in phylogenetic distance between disturbed and undisturbed environments should decrease with increasing elevation since the effect of reduced competition with disturbance is not as important at higher elevations.

## MATERIAL AND METHODS

### Sampling design and plant community sampling

#### Sampling design

Sampling was conducted along elevational gradients through a global standardized vegetation survey established in 2007 by the Mountain Invasion Research Network (MIREN, www.mountaininvasions.org; Haider et al. 2022). Survey data from 16 regions across the globe were included in this study (Fig. 2), with three mountain roads assessed in most regions, resulting in a total of 47 elevational gradients. The elevation gradients ranged from 0 to >4000 m a.s.l. and covered a wide range of climates and plant community types. Each elevational gradient was equally divided into 20 sample sites. At each sample site, two plots of 2 m x 50 m were surveyed: one roadside plot parallel and adjacent to the road (representing the disturbed plant community) and one natural plot perpendicular and 50 meters separated from the road (representing the less disturbed plant community), pointing to the centre of the first plot, with the midpoint at 75 m from the roadside plot (Figures 1a, b). Due to region-specific constraints (e.g., cliffs), not all sample sites could be realized (e.g., in Australia (Victoria) and Hawaii only roadside plots were established; see information for each region in Figure 2). In sum, we analysed native and non-native plants in 1157 sampling plots.

**Figure 2:**
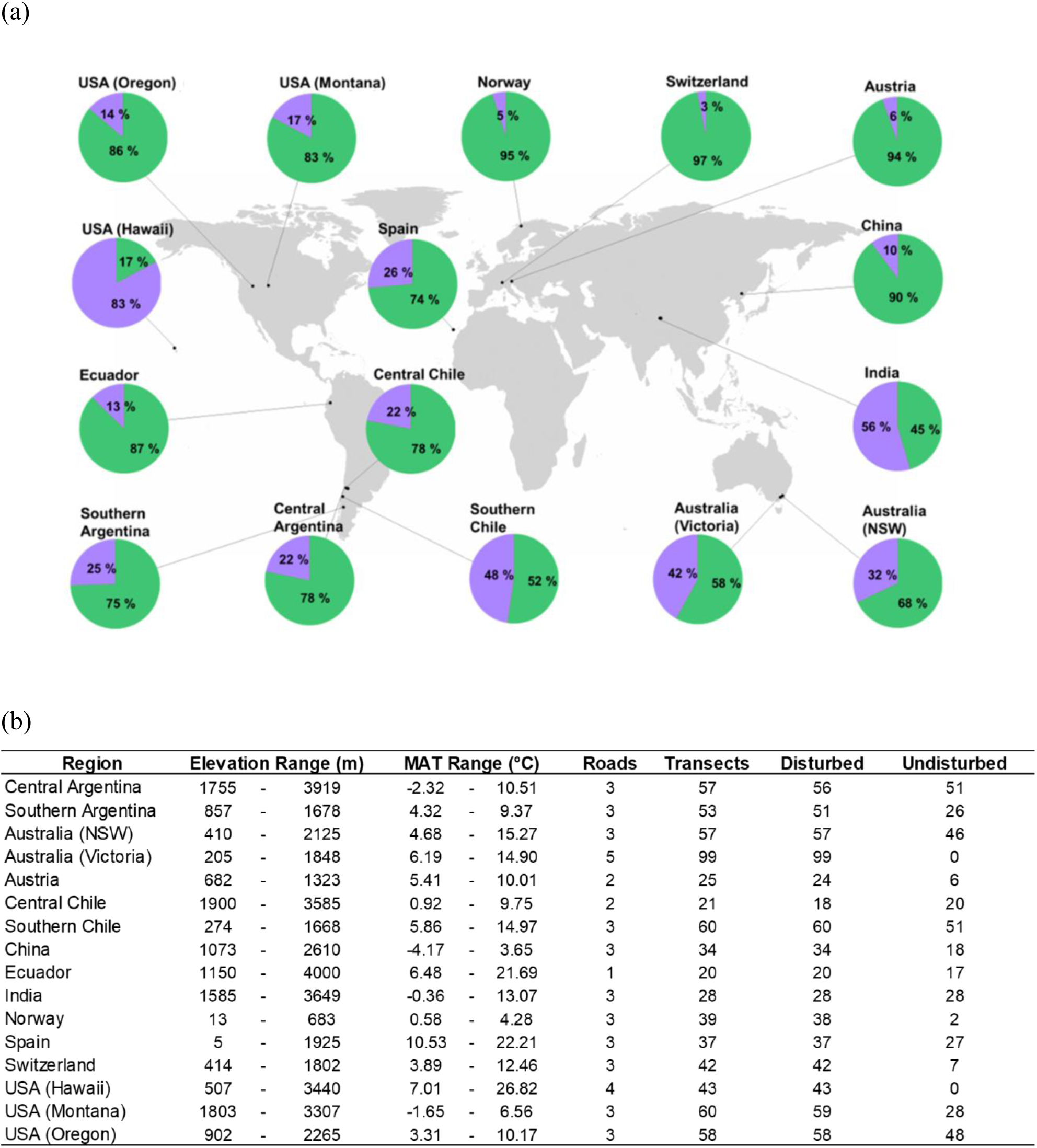
(a) The proportion of non-native (in purple) and native species (in green) for each region included in the study; (b) maximum and minimum elevation and mean annual temperature (MAT) for the study sites in each region, and the number of roads, sample sites, and disturbed and undisturbed plots for each region.

#### Plant community sampling

The presence of native and non-native vascular plant species was recorded in all plots following the MIREN protocol (Haider et al. 2022), thus ensuring a consistent spatial resolution. Total vegetation cover categories (1=< 0.1, 2=0.1 – 1, 3= 2 – 5, 4=6 – 10, 5= 11 – 25, 6= 26 – 50, 7= 51 – 75, 8= 76 – 100 percentage area) and abundance classes (1–10, 11–100, >100 individuals) of each species were visually estimated for each plot. Species names were harmonized between regions to detect potential synonyms and typographical errors using the R-packages “taxize” (Chamberlain et al. 2013), using “WorldFlora” (Kindt 2020) as the taxonomic backbone. Species names were matched with World Flora Online (http://www.worldfloraonline.org) and the Taxonomic Names Resolution Service (Boyle et al., 2013). The floristic status of the species (i.e., native or non-native to the region) was assigned based on regional floras and databases.

### Phylogeny and phylogenetic analysis

#### Phylogeny

We constructed a phylogenetic tree for all species recorded in all plots using the phylo.maker function in the ‘V.PhyloMaker’(Jin and Qian 2019) and ‘ape’ (Paradis et al. 2004) R-packages by matching the family, genus and species epithet from our survey with those in the backbone using the GBOTB.extended phylogeny in the ‘V.PhyloMaker’ R package (Jin and Qian 2022). All 190 families recorded in our survey, 1189 out of 1277 genera (> 93%) and 2261 of the total 3975 species (57%) were resolved in the mega-phylogeny. The number of unresolved species is not surprising, since our dataset includes a high number of endemic and rare species that have not been sequenced. We added absent genera and species to our phylogeny using the approach implemented in scenario three of V.PhyloMaker (Jin and Qian 2019), which is a robust method, commonly used in community phylogenetics (Dyderski and Jagodyinski 2020, Qian et al 2019). All analyses were performed in R version 4.02 (R Core Team, 2021).

#### Calculation of phylogenetic distances of non-native species to the native species in the community

To quantify the phylogenetic distance between each of the non-native species and the native community, we calculated four phylogenetic dissimilarity metrics suggested by Thuiller et al. 2010. For each non-native species in each sampling plot, we calculated the weighted and the unweighted mean distance to all native species (WMDNS and MDNS, respectively; Thuiller et al. 2010), the distance to the most abundant native species (DMANS), and the distance to its nearest native species (DNNS; both described in Thuiller et al. 2010). MDNS assumes that the entire community equally drives the establishment success of a non-native plant, while the WMDNS weights the MDNS by the abundances of the native species and assumes that the contribution of each native species depends on its relative abundance. All phylogenetic distance calculations were performed using the “ape” (Paradis et al. 2004) and “picante” packages (Kembel et al. 2010) and using the same code as in Thuiller et al. 2010 kindly provided by the first author. For each metric, we calculated the mean value across all non-native species per plot.

#### Phylogenetic structure of the native communities

To account for effects of different phylogenetic structures of the native communities influencing patterns of relatedness between non-native and native species, we calculated the mean phylogenetic distance among the native species, using the mean pairwise distance (MPD, Webb et al. 2002), and the standardized effect sizes of this metric (sMPD), using the mpd and the ses.mpd function from the R package ‘picante’ (Kembel et al. 2010). Positive values of sMPD and high p-values (> 0.95) indicate phylogenetic evenness (i.e., species within the community are more distantly related than expected by chance); negative values of sMPD and low p-values (< 0.05) indicate phylogenetic clustering (i.e., species within the community are more closely related than expected by chance).

### Statistical analyses: correlations between phylogenetic metrics and environmental factors

#### Global and regional patterns of phylogenetic relatedness of non-native to native species across elevation gradients

For our global analysis, we analyzed the effect of elevation and disturbance on the phylogenetic relatedness between non-native species and the native community using linear mixed-effects models for each of the four dissimilarity metrics (‘lmer’ in the R package ‘lmerTest’; Kuznetsova et al. 2015). Elevation was scaled between 0 and 1 in every region to allow comparison among regions with different elevational extents. For scaling, we used the ‘rescale’ function in the R package ‘scales’ (Wickham and Seidel 2022). Comparing elevational gradients across a wide spectrum of climatic zones offers an ideal system for testing hypotheses explaining ecological processes since increasing elevation is not only associated with variability in abiotic factors as decreasing temperature, but also with variability in biotic interactions and decreases of species diversity (Körner 2007). We included each dissimilarity metric as the response variable, and elevation, disturbance (disturbed or undisturbed plots) and their interaction as fixed effects. We included total vegetation cover as a covariate, to account for differences in the overall strength of competition. Sample site nested within road, nested within region were included as random effects to account for the nested structure of the sampling design. We fit each model twice, first with the linear term of elevation and second with its second-order polynomial to account for potential unimodal patterns of phylogenetic distance along the elevational gradient. For the second-order polynomial, we summarized the linear and the quadratic terms of elevation with the ‘poly’ function in R package stats, which uses QR factorization to generate monic orthogonal polynomials. We compared the linear and the quadratic fits based on the corrected Akaike information criterion (AICc) and selected the model with lowest AICc. Prior to analyses, we square-root transformed the response variables to satisfy the model assumptions of normal distribution.

To assess differences among regions, we fit additional models separately for each region. For the regional analyses, we only used WMDNS as the response variable, because results for the other metrics were similar (Table S1). We square-root transformed WMDNS to fulfil normal distribution assumptions. For each region, we compared models using a linear or quadratic of elevation, and selected the model with lower AICc.

#### Regional randomization models

We used randomizations to test whether non-native species are more or less phylogenetically related to the native species in each community than would be expected by chance. To do so, for each region, we generated a null distribution of non-native species within each community by randomizing the region’s non-native species in the phylogenetic distance matrix 1000 times using TaxaShuffle in the “picante” package in R (Kembel et al. 2010). Essentially, we generated 1000 communities with a random set of non-natives from the regional non-native species pool. We held the number of non-native species in each plot as well as the composition of the native community constant. For each randomized matrix, we calculated the phylogenetic distance of non-native to native species (using WMDNS) and ran the regional mixed models with interacting effects of elevation and disturbance (disturbed/undisturbed) as described above. For each randomized regression we extracted the fitted values and calculated their 95% confidence intervals. We compared observed model fits with the 95%CI of model fits for the random communities for each plot of each region to assess significance.

#### Global and regional patterns of phylogenetic structure of the native community across environmental gradients

Finally, we also ran the aforementioned global and regional linear mixed-effects models to test for the effect of elevation, disturbance and their interaction on the phylogenetic structure of the native community using mpd and sMPD as response variables. The results from these analyses can help to interpret how the phylogenetic structure of the native community contributes to the patterns of relatedness of non-native to native species. In the case of the native community, community assemblage and consequent patterns of phylogenetic relatedness could be determined not only by contemporary processes (like environmental filtering or competition) but also by the macroevolutionary history of diversification of the flora.

## RESULTS

### Global and regional patterns of phylogenetic relatedness of non-native to native species across environmental gradients

Since patterns of phylogenetic distance between non-native species and the resident native community were very similar for the four metrics used (see Table S1), we only present here the results for the weighted mean distance of the non-native species to all native species (WMDNS). At the global scale, the phylogenetic distance between non-native and native species declined with elevation, with steeper declines at higher elevation (significant quadratic effect of elevation; Table S2). This was also the case for some of the individual regions, while for other regions we found a linear negative relationship between phylogenetic distance and elevation (Table S2). Deviations from the global decrease were found for Ecuador and Spain (hump-shaped pattern) and for China and Switzerland (no elevational effects). At the global scale, we found phylogenetic distance of non-native to native species to be higher in roadside than in natural communities. This pattern was consistently observed in eight regions (Central and Southern Argentina, Australia (New South Wales), Southern Chile, Ecuador, Spain and USA (Montana and Oregon)), while there was no disturbance effect for the remaining eight regions (Figures 3 and 4, Tables S3 and S4). At the global scale, the slope of the negative relationship between elevation and phylogenetic distance was steeper for the roadside communities compared to natural ones (Figure 3, Table S3). At low and mid elevations, phylogenetic distance was greater in the roadside than in the natural communities. At highest elevations, the opposite pattern was found with greater phylogenetic distance in the natural communities. Regionally, we found the same pattern for Central and Southern Argentina and USA (Oregon), while we did not observe an interacting effect of elevation and disturbance for the other regions (Figure 4, Table S4).

**Figure 3.**
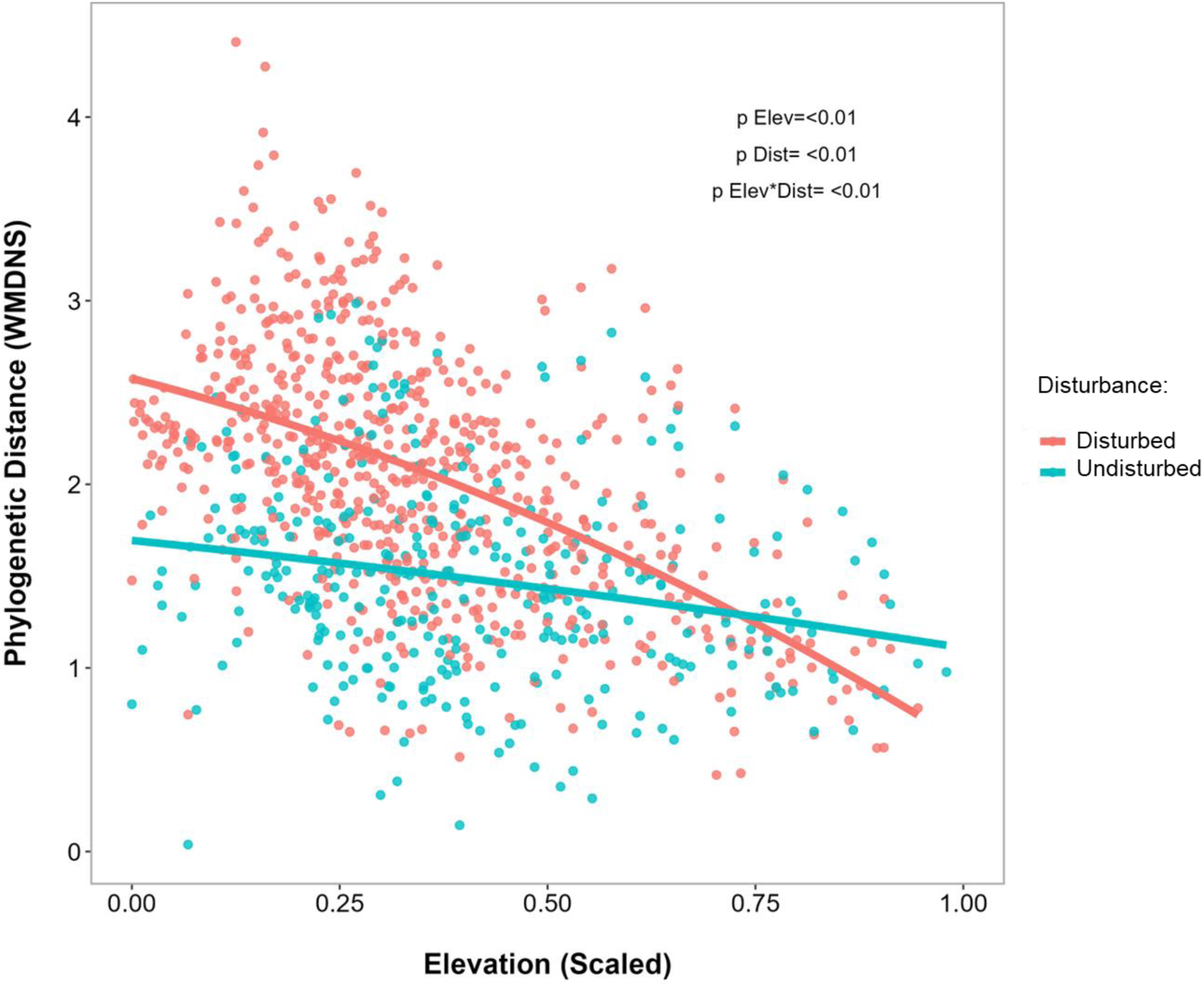
Global patterns of variation with elevation for Weighted Mean Phylogenetic Distance from the non-native to all native species (WMDNS; square root transformed) in roadsides (disturbed in the figure; red dots and lines) and natural plots (undisturbed in the figure; blue dots and lines). Curves of phylogenetic distance are based on model predictions. Note that elevation was scaled prior to modelling. Associated p-values (p) are shown for the effect of elevation (p Elev), disturbance (p Dist) and for the effect of the interaction between elevation and disturbance (p Elev*Dist).

**Figure 4.**
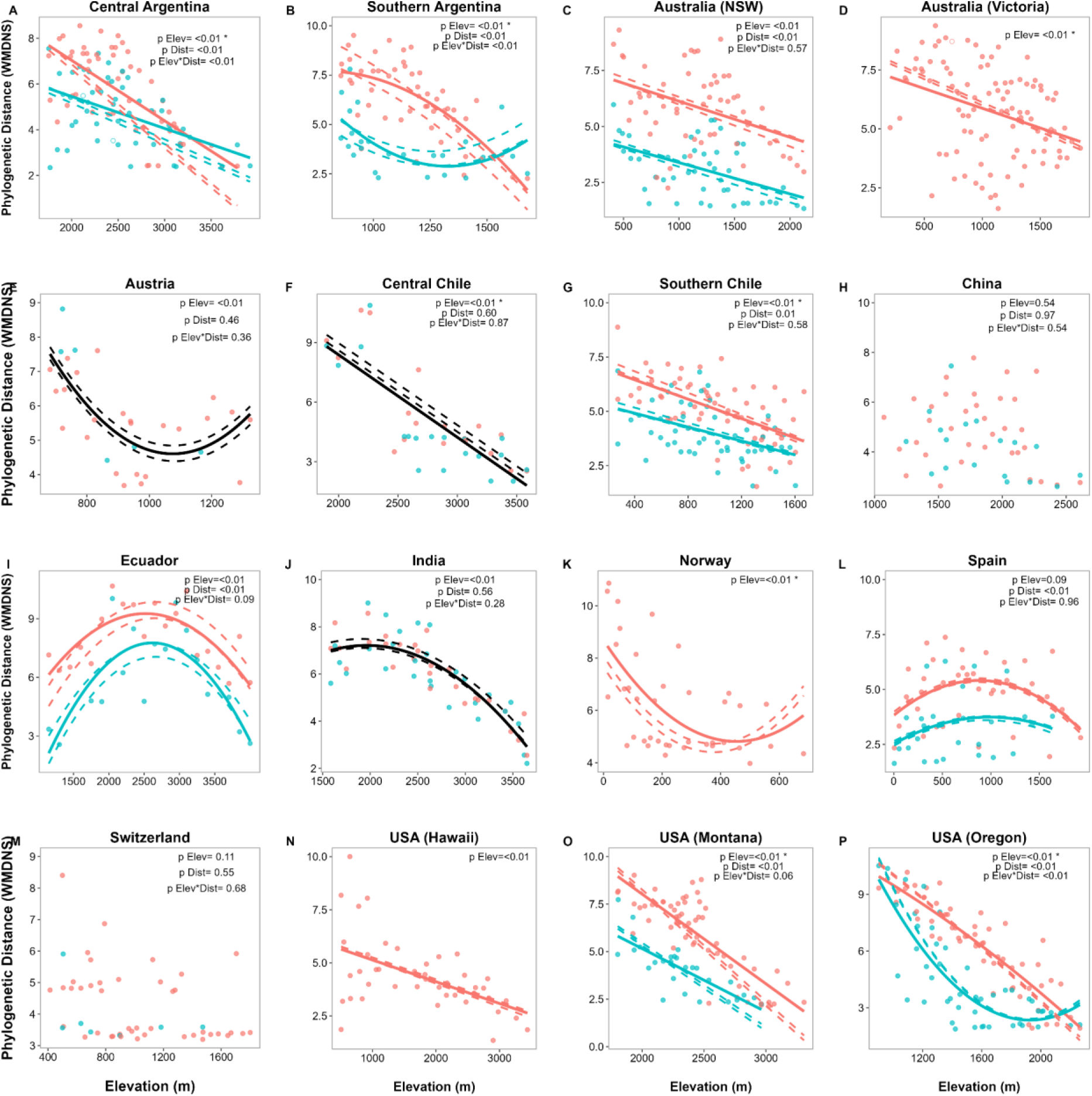
Regional patterns of variation with elevation for Weighted Mean Distance from the non-native species to all Native Species (WMDNS; sqrt-transformed) in disturbed roadside plots (red dots and lines) and undisturbed natural plots (blue dots and lines). Lines and curves of phylogenetic distance are based on model predictions (see Methods and Table S3). Black lines indicate that the effect of elevation was the same for roadside and natural plots (i.e. no effect of disturbance or the interaction between elevation and disturbance). If elevation, disturbance or their interaction was not significant, no lines were drawn. Solid lines represent the regression lines for the observed WMDNS values, while dotted lines represent the 0.5 and 95% confidence intervals for the predicted WMDNS values calculated with the randomized regressions. Each dot represents the observed WMDNS value for a specific plot, unfilled dots indicate that observed values were not significantly different than the predicted values (based on a t.test comparison). Associated p-values (p) are shown for each model predictor for the effect of elevation (p elevation) and for the effect of the interaction between elevation and disturbance (p interaction). * Indicates when observed estimates were not within the confidence intervals of the predicted values of the randomized models.

Across all regions, we found an effect of total vegetation cover on the phylogenetic distance of non-native species to the native community, whereby non-native species were more distantly related to the native community with increasing total vegetation cover (Figure S1, Table S3).

Whitin regions, we found a significant positive effect of cover on phylogenetic distance for Central Argentina, Australia (Victoria) and USA (Hawaii) but a negative effect for Southern Chile (Figure S2, Table S4).

### Regional randomization models

For eight of the 12 regions for which we found a significant negative correlation between elevation and phylogenetic distance, the observed effect was different from the relationship in randomized communities (i.e. the observed estimates were not within the confidence intervals of the predicted values of the randomization models; Figure 4). However, the comparison of the observed pattern with the randomized regression varied between regions, with observed phylogenetic distances greater or smaller than the predicted ones along the whole elevational gradient, at specific elevations only, or either for roadside or natural communities only (Figure 4). Specifically, we found smaller phylogenetic distances than expected by chance for the roadside communities in Australia (Victoria; towards lower elevations) and Norway (towards higher elevations), along the entire gradient for central Chile, and for roadside and natural low elevation communities in southern Chile and USA (Montana and Oregon). Contrarily, we found greater phylogenetic distance than expected by chance for roadside and natural communities in central Argentina (all elevations) and USA (Montana, towards higher elevations), for roadside communities towards higher elevations in southern Argentina, Australia (Victoria) and USA (Oregon), as well as for roadside communities towards lower elevations in Ecuador and Norway.

### Global and regional patterns of the phylogenetic structure of native communities across environmental gradients

Both across all regions and within most regions, the native communities were more phylogenetically clustered than expected by chance with no significant effect of elevation (Table S5, Figure S3). We did find an effect of disturbance on the phylogenetic structure of native communities, with roadside communities displaying significantly more phylogenetic clustering than natural communities. This effect did not change with elevation (Table S5 and S6, Figure S3 and S4).

## DISCUSSION

Our study revisited Darwin’s Naturalization Conundrum by exploring how patterns of relatedness between non-native and native plant species vary along elevation and disturbance gradients, and by seeking insights into the role of ecological processes in generating these patterns. We found support for our first hypothesis, that with increasing elevation non-native species were more closely related to the native community, suggesting that pre-adaptation is an important determinant of invasion success in stressful habitats. However, contrary to our second and third hypotheses, non-native plants were less closely related to the native communities in disturbed environments (i.e., roadsides) compared to natural habitats and the increase in relatedness with elevation was more pronounced for anthropogenically disturbed sites. This suggests that competition might be promoting similar species by reducing fitness differences in more competitive communities (Mayfield & Levine 2010) or that disturbance may promote the establishment of non-native plants through mechanisms other than competition filters, such as opening of new niches (Catford et al. 2012, Moles et al. 2008). Our findings are consistent with the existence of different environmental filters during the invasion process as described by Dietz and Edwards (2006) and observed for invasions in mountain systems by McDougall et al. (2018). Accordingly, non-native species are first introduced in lowland environments and then spread in two directions: i) vertical dispersal to higher elevations along roadside corridors (Alexander 2011; Haider et al. 2018), and ii) horizontal dispersal to the adjacent less disturbed natural habitats (McDougall et al. 2018).

### First invasion filter: vertical dispersion towards higher elevations along roads

Non-native species enter mountain systems at low elevations and from there they spread upwards, often using roads as dispersal corridors (Alexander et al. 2011, Arevalo et al. 2010). During this vertical spread, non-natives face changes in the abiotic environmental conditions, and particularly high elevations are commonly associated with more stressful conditions (e.g. via decreasing temperatures). In contrast to non-native species being in the phase of spread after establishment, native species had thousands and millions of years to spread across the landscape and patterns of relatedness may be also influenced by their macroevolutionary history of dispersion. Our observation that non-native become closer related to native species with increasing elevation supports the hypothesis that environmental filtering is filtering-out non-native species that are poorly adapted to the new abiotic conditions that they encounter as they spread upwards along the gradient (Cadotte and Tucker 2017). Consistent with our findings, two recent studies report that non-native plant species are more closely related to native species in harsher climates, indicating a greater influence of preadaptation. More specifically, Fan et al. (2023) found a closer phylogenetic relationship between non-native and native species at higher than at lower latitudes analysing a global database of naturalized non-native plants, and Wang et al. (2024) found that the invasive plant *Ageratina adenophora* was more successful at invading closely related plant communities in low temperature, high elevation habitats in a mountain region of China.

While we found that non-native and native species become closer related with elevation for roadside and natural environments, at low and mid-elevations non-native and native plants were more distantly related in disturbed roadsides than in less disturbed natural habitats, which is contrary to the assumption that reduced competition in disturbed habitats lifts the burden of limiting similarity as a requirement for coexistence (Götzenberger et al. 2012). Two characteristics of disturbed roadside environments may promote the entrance particularly of more distantly related non-native species: the high propagule pressure and the opening of new available niches. Mountain roads constitute a gradient in propagule pressure, which is an overriding factor facilitating invasion (Fuentes-Lillo et al. 2021, Catford et al. 2011, Holle and Simberloff 2005, Fine 2004). Due to anthropogenic activities, lowlands are the main source of propagules of non-native species and provide a large pool of potential invaders spreading to higher elevations (Pauchard and Alaback 2004; Lembrechts et al. 2014). The non-native species pool of low elevations may be particularly diverse because of a high density of settlements and cities together with an intentional introduction of functionally (and hence typically phylogenetically) dissimilar species for example for ornamental purposes (like exotic species collections in public or private gardens; (Padullés Cubino et al. 2019)) might increase the phylogenetic distance between non-native and native species. The high propagule pressure in the lowland roadside environments and the availability of new niches opened through the road disturbance might favour the establishment of non-native species that are phylogenetically distinct to the native assemblages. When non-native species continue dispersing upwards along roads, the effect of propagule pressure decreases and the effect of environmental filters increases. In addition to the effect of propagule pressure and new niches, Clavel et al. (2023) found that roadside disturbance promotes plant species associated with arbuscular mycorrhizal (AM), which have been found to be phylogenetically more distinct than ectomycorrhizal- or ericoid-mycorrhizal- associated species, and that non-native plant species (usually AM plants) are more successful in environments dominated by AM associations.

### <u>Second invasion filter: horizontal dispersion from disturbed roadsides to adjacent less</u> <u>disturbed natural communities</u>

Once established in roadside environments, some non-native species may undergo a secondary invasion (Dietz and Edwards 2006) and spread away from the road into the adjacent vegetation after passing a second environmental filter related to different habitat conditions in natural environments, such as differences in light or soil moisture (Cavender-Bares et al. 2009, McDougall et al. 2018). The subset of non-native species being able establish away from the roadside are the ones potentially better preadapted to the new conditions and hence more closely related to the native species in the less disturbed natural environments. In addition to different abiotic conditions, natural environments adjacent to roads present also changes in biotic components such as differences in competition or mutualistic interactions. Further restrictions may take place for non-native specieś establishment through biotic competition with the already established species. To cope with more competitive natural environments, non-native species need to have sufficient phylogenetic distance to native species to overcome biotic competition and be able to use empty niches. This could explain why phylogenetic distance from non-native to native species was always greater at low than at high elevations, as competition is generally lower at high elevations (Callaway et al. 2002). Our finding that with increasing vegetation cover (more competitive communities) phylogenetic distance from the non-native to the native species also increases supports the assumption that competition leads to higher phylogenetic distance in natural environments. Our results align with a study by Omer et al. (2022) showing that environmental filtering is important for the naturalization of non-native species, but biotic competition is important for them to spread and become more dominant in the community.

### From patterns of phylogenetic distance to ecological processes

Our results show for most regions that the observed variation in patterns of phylogenetic distance along gradients differs from random expectations. This strengthens the interpretation that patterns of relatedness from non-native to native species are driven by ecological processes and are not simply the result of random assembly. The decrease in phylogenetic distance as non-native species spread upward and into natural communities points towards an effect of environmental filtering favoring preadapted species. This interpretation is strengthened by the randomized regional models when observed patterns of phylogenetic distance were smaller than expected by chance. In more competitive natural environments, some regions showed higher phylogenetic distances than expected by chance, suggesting an effect of competition leading to limiting similarity. However, other regions showed smaller phylogenetic distances than expected by chance, which could indicate a filtering by competition that favors more competitive species. Observing greater phylogenetic distance than expected by chance in roadside communities reinforces the role of processes such as the creation of new niches through disturbance and a larger non-native species pool favoring more diverse species within the individual communities. In addition, we showed that the observed patterns of variation in phylogenetic relatedness between non-native species and the native community are not driven by non-contemporary factors associated to the assemblage of the native communities as a result of their macroevolutionary history. The general clustering pattern of the native communities indicates that the pattern of greater phylogenetic distance of non-native species to the native community towards low elevations is not driven by a pattern of overdispersion of the native community but supports our assumption of environmental filtering along the elevational gradient which selects non-native species preadapted to the environment.

Variability between regions might be explained by region-specific situations, as differences in disturbance intensity. For example, a low level of disturbance in native communities might align with higher biotic resistance against the establishment of non-native species. Region-specific differences might also arise from differences in the species pool entering each region based on socio-economic and historic factors and regulations (Aronson et al. 2016). In addition, other ecological processes such as antagonistic or mutualistic interactions (with important effects at the community scale) may influence patterns of relatedness between native and non-native species. For example, closely related species are more likely to share pathogens, parasites, and herbivores than distantly related species (Ness et al. 2011; Ødegaard et al. 2005). Shared enemies may act as a biotic filter for non-native species closely related to the native community. There remains a need to examine changes in the functional dynamics of plant communities as a result of plant invasions, and how these changes alter the ecological interactions of plants with other organisms, in order to fully understand the factors that determine the success of non-native species and the consequences of invasions.

Overall, we show that habitats at lower elevations and with higher levels of disturbance are characterized by greater phylogenetic distances of non-native species to the native community. Our results suggest that different environmental filters are the main drivers of non-native species distributions in mountain environments. Roadside disturbance had a strong selective effect on the native community, while non-native species occurrence seems to depend strongly on propagule pressure, particularly at low elevations. Elevation-related filtering (e.g. climate filtering) restricts the upward spread of non-native species, while other filters related to changes in habitat conditions hinders non-native species from percolating into natural adjacent vegetation. Still, competition seems to play a role for non-native species establishment, particularly in less disturbed habitats, with decreasing importance with increasing elevation. We conclude that both the pre-adaptation hypothesis and Darwin’s naturalization hypothesis are valid when applied to the appropriate environmental conditions.

Finally, our results highlight important implications of plant invasions under ongoing global change. The occurrence of plant invasions is predicted to increase under global change (Liu et al. 2017). Increasing temperatures and habitat disturbance may reduce the effect of environmental filters that filter out phylogenetically distinct species. Therefore, more non-native species may reach habitats with a lower presence of invasions, such as the alpine environment (Lembrechts et al. 2016). In addition, another consequence is that, assuming that phylogenetic distance is associated with trait functional dissimilarity (Webb 2000), increasing habitat disturbance and rising global temperatures could favor non-native invasive species that not only change the species composition of the community but also alter ecosystem functions.

## ACKNOWLEDGMENTS

We thank all the members of the Mountain Invasion Research Network (https://www.mountaininvasions.org/) who helped to collect the data that made this study possible. We thank Wilfried Thuiller for providing the R code to compute the phylogenetic distance metrics. We thank Helge Bruelheide and Matthias Grenié for their help with the coding. We also thank William C. Wetzel for his kind comments on this manuscript. This research was funded by the German Research Foundation for support via the Flexpool Funding program of the German Centre for Integrative Biodiversity Research (iDiv) – Halle, Jena, Leipzig. LAC, EF-L and AP acknowledge funding by Fondecyt 12131616, 1211197, ART 210038 and ANID/BASAL FB210006, and MAS acknowledges the financial support by the Ministry of Education, Govt. of India & Mission Directorate RUSA J&K under RUSA 2.0 Project.

## Notes

### Competing Interest Statement

The authors have declared no competing interest.

## REFERENCES

1. Alexander, J. M., Kueffer, C., Daehler, C. C., Edwards, P. J., Pauchard, A., Seipel, T., … & Walsh, N. (2011). Assembly of nonnative floras along elevational gradients explained by directional ecological filtering. Proceedings of the National Academy of Sciences, 108(2), 656–661.

2. Arevalo JR, Otto R, Escudero C, Fernandez-Lugo S, Arteaga M, Delgado JD & Fernandez-Palacios JM. 2010. Do anthropogenic corridors homogenize plant communities at a local scale? A case studied in Tenerife (Canary Islands). Plant Ecology 209: 23–35. doi:10.1007/s11258-009-9716-y.

3. Aronson, M. F., Nilon, C. H., Lepczyk, C. A., Parker, T. S., Warren, P. S., Cilliers, S. S., … & Zipperer, W. (2016). Hierarchical filters determine community assembly of urban species pools. Ecology, 97(11), 2952–2963.

4. Bellard, C., Marino, C., & Courchamp, F. (2022). Ranking threats to biodiversity and why it doesn’t matter. Nature Communications, 13(1), 1–4.

5. Boyle, B., Hopkins, N., Lu, Z., Raygoza Garay, J. A., Mozzherin, D., Rees, T., … & Enquist, B. J. (2013). The taxonomic name resolution service: an online tool for automated standardization of plant names. BMC bioinformatics, 14, 1–15.

6. Cadotte, M. W., Campbell, S. E., Li, S. P., Sodhi, D. S., & Mandrak, N. E. (2018). Preadaptation and naturalization of nonnative species: Darwin’s two fundamental insights into species invasion. Annual Review of Plant Biology, 661-684.

7. Cadotte, M. W., & Tucker, C. M. (2017). Should environmental filtering be abandoned?. Trends in Ecology & Evolution, 32(6), 429–437.

8. Callaway, et al., Positive interactions among alpine plants increase with stress. Nature 417, 844–848 (2002)

9. Carboni, M., A. T. R. Acosta, and C. Ricotta. 2013. Are differences in functional diversity among plant communities on Mediterranean coastal dunes driven by their phylogenetic history? Journal of Vegetation Science 24:932–941.

10. Catford, J. A., C. C. Daehler, H. T. Murphy, A. W. Sheppard, B. D. Hardesty, D. A. Westcott, M. Rejmánek, P. J. Bellingham, J. Pergl, and C. C. Horvitz. 2012. The intermediate disturbance hypothesis and plant invasions: Implications for species richness and management. Perspectives in Plant Ecology, Evolution and Systematics 14:231–241.

11. Catford, J. A., Vesk, P. A., White, M. D., & Wintle, B. A. (2011). Hotspots of plant invasion predicted by propagule pressure and ecosystem characteristics. Diversity and Distributions, 17(6), 1099–1110.

12. Cavender-Bares, J., Ackerly, D. D., Baum, D. A., & Bazzaz, F. A. (2004). Phylogenetic overdispersion in Floridian oak communities. The American Naturalist, 163(6), 823–843.

13. Cavender-Bares, J., A. Keen, and B. Miles. 2006. Phylogenetic structure of Floridian plant communities depends on taxonomic and spatial scale. Ecology 87:S109–S122.

14. Cavender-Bares, J., Kozak, K. H., Fine, P. V., & Kembel, S. W. (2009). The merging of community ecology and phylogenetic biology. Ecology Letters, 12(7), 693–715.

15. Chamberlain, S. A., & Szöcs, E. (2013). taxize: taxonomic search and retrieval in R. F1000Research, 2.

16. Clavel, J., Lembrechts, J. J., Lenoir, J., Haider, S., McDougall, K., Nuñez, M. A., … & Nijs, I. (2024). Roadside disturbance promotes plant communities with arbuscular mycorrhizal associations in mountain regions worldwide. Ecography, e07051.

17. Daehler, C. C. 2001. Darwin’s naturalization hypothesis revisited. The American Naturalist 158:324–330.

18. Dietz, H., & Edwards, P. J. (2006). Recognition that causal processes change during plant invasion helps explain conflicts in evidence. Ecology, 87(6), 1359–1367.

19. Diez, J. M., Sullivan, J. J., Hulme, P. E., Edwards, G., & Duncan, R. P. (2008). Darwin’s naturalization conundrum: dissecting taxonomic patterns of species invasions. Ecology Letters, 11(7), 674–681.

20. Dyderski, M. K., & Jagodziński, A. M. (2020). Impact of invasive tree species on natural regeneration species composition, diversity, and density. Forests, 11(4), 456.

21. Faith, D. P. (1992). Conservation evaluation and phylogenetic diversity. Biological Conservation, 61(1), 1–10.

22. Fan, S. Y., Yang, Q., Li, S. P., Fristoe, T. S., Cadotte, M. W., Essl, F., … & van Kleunen, M. (2023). A latitudinal gradient in Darwin’s naturalization conundrum at the global scale for flowering plants. Nature Communications, 14(1), 6244.

23. Fuentes-Lillo, E., Lembrechts, J. J., Cavieres, L. A., Jiménez, A., Haider, S., Barros, A., & Pauchard, A. (2021). Anthropogenic factors overrule local abiotic variables in determining non-native plant invasions in mountains. Biological Invasions, 23(12), 3671–3686.

24. Gerhold, P., Cahill Jr, J. F., Winter, M., Bartish, I. V., & Prinzing, A. (2015). Phylogenetic patterns are not proxies of community assembly mechanisms (they are far better). Functional Ecology, 29(5), 600–614.

25. Gerhold, P., Pärtel, M., Tackenberg, O., Hennekens, S. M., Bartish, I., Schaminée, J. H., … & Prinzing, A. (2011). Phylogenetically poor plant communities receive more alien species, which more easily coexist with natives. The American Naturalist, 177(5), 668–680.

26. Godoy, O., Kraft, N. J., & Levine, J. M. (2014). Phylogenetic relatedness and the determinants of competitive outcomes. Ecology Letters, 17(7), 836–844.

27. Götzenberger, L., de Bello, F., Bråthen, K. A., Davison, J., Dubuis, A., Guisan, A., … & Zobel, M. (2012). Ecological assembly rules in plant communities—approaches, patterns and prospects. Biological reviews, 87(1), 111–127.

28. Güsewell, S., & Klötzli, F. (2012). Local plant species replace initially sown species on roadsides in the Swiss National Park. eco. mont: Journal on Protected Mountain Areas Research and Management, 4(1), 23-34.

29. Hadley Wickham and Dana Seidel (2022). scales: Scale Functions for Visualization. R package version 1.2.0. https://CRAN.R-project.org/package=scales

30. Haider, S., J. Lembrechts, K. McDougall, A. Pauchard, J. M. Alexander, A. Barros, L. Cavieres, I. Rashid, L. Rew, and A. Aleksanyan. 2021. Think globally, measure locally: The MIREN standardized protocol for monitoring species distributions along elevation gradients. Authorea Preprints.

31. Holle, B. V., & Simberloff, D. (2005). Ecological resistance to biological invasion overwhelmed by propagule pressure. Ecology, 86(12), 3212–3218.

32. Hughes, C., and R. Eastwood. 2006. Island radiation on a continental scale: exceptional rates of plant diversification after uplift of the Andes. Proceedings of the National Academy of Sciences 103:10334–10339.

33. IPBES (2023). Summary for Policymakers of the Thematic Assessment Report on Invasive Alien Species and their Control of the Intergovernmental Science-Policy Platform on Biodiversity and Ecosystem Services.

34. Roy, H. E., Pauchard, A., Stoett, P., Renard Truong, T., Bacher, S., Galil, B. S., Hulme, P. E., Ikeda, T., Sankaran, K. V., McGeoch, M. A., Meyerson, L. A., Nuñez, M. A., Ordonez, A., Rahlao, S. J., Schwindt, E., Seebens, H., Sheppard, A. W., and Vandvik, V. (eds.). IPBES secretariat, Bonn, Germany. 10.5281/zenodo.7430692

35. Jiang, L., J. Tan, and Z. Pu. 2010. An experimental test of Darwin’s naturalization hypothesis. The American Naturalist 175:415–423.

36. Jin, Y., and H. Qian. 2019. V. PhyloMaker: an R package that can generate very large phylogenies for vascular plants. Ecography 42:1353–1359.

37. Jin, Yi, and Hong Qian. “V. PhyloMaker2: An updated and enlarged R package that can generate very large phylogenies for vascular plants.” Plant Diversity 44.4 (2022): 335- 339.

38. Jones, E. I., S. L. Nuismer, and R. Gomulkiewicz. 2013. Revisiting Darwin’s conundrum reveals a twist on the relationship between phylogenetic distance and invasibility. Proceedings of the National Academy of Sciences 110:20627–20632.

39. Karger, D.N., Conrad, O., Böhner, J., Kawohl, T., Kreft, H., Soria-Auza, R.W., Zimmermann, N.E., Linder, P., Kessler, M. (2017): Climatologies at high resolution for the Earth land surface areas. Scientific Data. 4

40. Kembel, S. W., Cowan, P. D., Helmus, M. R., Cornwell, W. K., Morlon, H., Ackerly, D. D., … & Webb, C. O. (2010). Picante: R tools for integrating phylogenies and ecology. Bioinformatics, 26(11), 1463–1464.

41. Kindt, R. 2020. WorldFlora: An R package for exact and fuzzy matching of plant names against the World Flora Online taxonomic backbone data. Applications in Plant Sciences 8.

42. Körner, C., & Life, A. P. (2003). Functional plant ecology of high mountain ecosystems. Alpine plant life, 1-7.

43. Körner, C. 2007. The use of ’altitude’ in ecological research. Trends in Ecology & Evolution 22:569–574.

44. Kraft, N. J., Adler, P. B., Godoy, O., James, E. C., Fuller, S., & Levine, J. M. (2015). Community assembly, coexistence and the environmental filtering metaphor. Functional ecology, 29(5), 592–599.

45. Kuznetsova, A., Brockhoff, P. B., & Christensen, R. H. B. (2015). Package ‘lmertest’. R package version, 2(0), 734.

46. Lembrechts, J. J., Milbau, A., & Nijs, I. (2014). Alien roadside species more easily invade alpine than lowland plant communities in a subarctic mountain ecosystem. PloS one, 9(2), e89664.

47. Lembrechts, J. J., Pauchard, A., Lenoir, J., Nuñez, M. A., Geron, C., Ven, A., … & Milbau, A. (2016). Disturbance is the key to plant invasions in cold environments. Proceedings of the National Academy of Sciences, 113(49), 14061–14066.

48. Li, D., L. Trotta, H. E. Marx, J. M. Allen, M. Sun, D. E. Soltis, P. S. Soltis, R. P. Guralnick, and B. Baiser. 2019. For common community phylogenetic analyses, go ahead and use synthesis phylogenies. Ecology 100:e02788.

49. Liu, Y., Oduor, A. M., Zhang, Z., Manea, A., Tooth, I. M., Leishman, M. R., … & Van Kleunen, M. (2017). Do invasive alien plants benefit more from global environmental change than native plants?. Global Change Biology, 23(8), 3363–3370.

50. Lososová, Z., de Bello, F., Chytrý, M., Kühn, I., Pyšek, P., Sádlo, J., … & Zelený, D. (2015). Alien plants invade more phylogenetically clustered community types and cause even stronger clustering. Global Ecology and Biogeography, 24(7), 786–794.

51. Ma, C., S. Li, Z. Pu, J. Tan, M. Liu, J. Zhou, H. Li, and L. Jiang. 2016. Different effects of invader–native phylogenetic relatedness on invasion success and impact: a meta- analysis of Darwin’s naturalization hypothesis. Proceedings of the Royal Society B: Biological Sciences 283:20160663.

52. Maitner, B. S., D. S. Park, B. J. Enquist, and K. M. Dlugosch. 2021. Where we’ve been and where we’re going: the importance of source communities in predicting establishment success from phylogenetic relationships. Ecography.

53. Malecore, E. M., W. Dawson, A. Kempel, G. Müller, and M. van Kleunen. 2019. Nonlinear effects of phylogenetic distance on early-stage establishment of experimentally introduced plants in grassland communities. Journal of Ecology 107:781–793.

54. Mayfield, M. M., & Levine, J. M. (2010). Opposing effects of competitive exclusion on the phylogenetic structure of communities. Ecology letters, 13(9), 1085–1093.

55. McDougall, K. L., Lembrechts, J., Rew, L. J., Haider, S., Cavieres, L. A., Kueffer, C., … & Alexander, J. M. (2018). Running off the road: roadside non-native plants invading mountain vegetation. Biological invasions, 20(12), 3461–3473.

56. McKinney, M. L. 2004. Measuring floristic homogenization by non-native plants in North America. Global Ecology and Biogeography 13:47–53.

57. Moles, A. T., Gruber, M. A., & Bonser, S. P. (2008). A new framework for predicting invasive plant species. Journal of Ecology, 96(1), 13–17.

58. Moles, A. T., Flores-Moreno, H., Bonser, S. P., Warton, D. I., Helm, A., Warman, L., … & Thomson, F. J. (2012). Invasions: the trail behind, the path ahead, and a test of a disturbing idea. Journal of Ecology, 100(1), 116–127.

59. Narwani, A., Matthews, B., Fox, J., & Venail, P. (2015). Using phylogenetics in community assembly and ecosystem functioning research. Functional Ecology, 29(5), 589–591.

60. Ness, J. H., Rollinson, E. J., & Whitney, K. D. (2011). Phylogenetic distance can predict susceptibility to attack by natural enemies. Oikos, 120(9), 1327–1334.

61. Omer, A., Fristoe, T., Yang, Q., Razanajatovo, M., Weigelt, P., Kreft, H., … & van Kleunen, M. (2022). The role of phylogenetic relatedness on alien plant success depends on the stage of invasion. Nature Plants, 1-9.

62. Ødegaard, F., Diserud, O. H., & Østbye, K. (2005). The importance of plant relatedness for host utilization among phytophagous insects. Ecology Letters, 8(6), 612–617.

63. Padullés Cubino, J., Cavender-Bares, J., Hobbie, S. E., Hall, S. J., Trammell, T. L., Neill, C., … & Groffman, P. M. (2019). Contribution of non-native plants to the phylogenetic homogenization of US yard floras. Ecosphere, 10(3), e02638.

64. Park, D. S., X. Feng, B. S. Maitner, K. C. Ernst, and B. J. Enquist. 2020. Darwin’s naturalization conundrum can be explained by spatial scale. Proceedings of the National Academy of Sciences 117:10904–10910.

65. Pauchard A, Kueffer C, Dietz H, Daehler CC, Alexander J, Edwards PJ, Arévalo JR, Cavieres L, Guisan A, Haider S, Jakobs G, McDougall KL, Millar CI, Naylor BJ, Parks CG, Rew LJ, Seipel T (2009) Ain’t no mountain high enough: plant invasions reaching high elevations. Front Ecol Environ 7:479–486

66. Peay, K. G., M. Belisle, and T. Fukami. 2012. Phylogenetic relatedness predicts priority effects in nectar yeast communities. Proceedings of the Royal Society B: Biological Sciences 279:749–758.

67. Petsch, D. K., Bertoncin, A. P. D. S., Ortega, J. C. G., & Thomaz, S. M. (2022). Non-native species drive biotic homogenization, but it depends on the realm, beta diversity facet and study design: a meta-analytic systematic review. Oikos.

68. Prinzing, A., C. A. D’Haese, S. Pavoine, and J. Ponge. 2014. Species living in harsh environments have low clade rank and are localized on former Laurasian continents: a case study of Willemia (Collembola). Journal of biogeography 41:353–365.

69. Procheş, Ş., J. R. U. Wilson, and R. M. Cowling. 2006. How much evolutionary history in a 10× 10 m plot? Proceedings of the Royal Society B: Biological Sciences 273:1143– 1148.

70. Procheş, Ş., J. R. U. Wilson, D. M. Richardson, and M. Rejmánek. 2008. Searching for phylogenetic pattern in biological invasions. Global Ecology and Biogeography 17:5– 10.

71. Pyšek, P., P. E. Hulme, D. Simberloff, S. Bacher, T. M. Blackburn, J. T. Carlton, W. Dawson, F. Essl, L. C. Foxcroft, and P. Genovesi. 2020. Scientists’ warning on invasive alien species. Biological Reviews 95:1511–1534.

72. Pyšek, P., V. Jarošík, P. E. Hulme, J. Pergl, M. Hejda, U. Schaffner, and M. Vilà. 2012. A global assessment of invasive plant impacts on resident species, communities and ecosystems: the interaction of impact measures, invading species’ traits and environment. Global Change Biology 18:1725–1737.

73. Qian, H., Deng, T., Jin, Y., Mao, L., Zhao, D., & Ricklefs, R. E. (2019). Phylogenetic dispersion and diversity in regional assemblages of seed plants in China. Proceedings of the National Academy of Sciences, 116(46), 23192–23201.

74. Qian, H., and Y. Jin. 2016. An updated megaphylogeny of plants, a tool for generating plant phylogenies and an analysis of phylogenetic community structure. Journal of Plant Ecology 9:233–239.

75. Qian, H., and Y. Jin. 2021. Are phylogenies resolved at the genus level appropriate for studies on phylogenetic structure of species assemblages? Plant Diversity 43:255–263.

76. Qian, H., Y. Jin, F. Leprieur, X. Wang, and T. Deng. 2020. Geographic patterns and environmental correlates of taxonomic and phylogenetic beta diversity for large-scale angiosperm assemblages in China. Ecography 43:1706–1716.

77. Qian, H., Y. Jin, F. Leprieur, X. Wang, and T. Deng. 2021. Patterns of phylogenetic beta diversity measured at deep evolutionary histories across geographical and ecological spaces for angiosperms in China. Journal of Biogeography 48:773–784.

78. Qian, H., & Sandel, B. (2022). Darwin’s preadaptation hypothesis and the phylogenetic structure of native and alien regional plant assemblages across North America. Global Ecology and Biogeography, 31(3), 531–545.

79. Qian, H., and J. Zhang. 2016. Are phylogenies derived from family-level supertrees robust for studies on macroecological patterns along environmental gradients? Journal of systematics and evolution 54:29–36.

80. R Core Team. (2021). R: A language and environment for statistical computing. R Foundation for Statistical Computing, Vienna, Austria. URL. https://www.R-project.org/.

81. Ratier Backes, A., Römermann, C., Alexander, J. M., Arévalo, J. R., Keil, P., Padrón- Mederos, M. A., … & Haider, S. (2023). Mechanisms behind elevational plant species richness patterns revealed by a trait-based approach. Journal of Vegetation Science, 34(1), e13171.

82. Rejmánek, M. 1996. A theory of seed plant invasiveness: the first sketch. Biological conservation 78:171–181.

83. Richardson, D. M., Pyšek, P., Rejmanek, M., Barbour, M. G., Panetta, F. D., & West, C. J. (2000). Naturalization and invasion of alien plants: concepts and definitions. Diversity and distributions, 6(2), 93–107.

84. Ricotta, C., F. A. la Sorte, P. Pyšek, G. L. Rapson, L. Celesti-Grapow, and K. Thompson. 2009. Phyloecology of urban alien floras. Journal of Ecology 97:1243–1251.

85. Smith, S. A., and J. W. Brown. 2018. Constructing a broadly inclusive seed plant phylogeny. American journal of botany 105:302–314.

86. Stubbs, W.J. & Wilson, J.B. (2004) Evidence for limiting similarity in a sand dune community. Journal of Ecology, 92, 557–567

87. Swenson, N. G., & Enquist, B. J. (2007). Ecological and evolutionary determinants of a key plant functional trait: wood density and its community-wide variation across latitude and elevation. American journal of botany, 94(3), 451–459.

88. Swenson, N. G. 2013. The assembly of tropical tree communities–the advances and shortcomings of phylogenetic and functional trait analyses. Ecography 36:264–276.

89. Thuiller, W., L. Gallien, I. Boulangeat, F. de Bello, T. Münkemüller, C. Roquet, and S. Lavergne. 2010. Resolving Darwin’s naturalization conundrum: a quest for evidence. Diversity and distributions 16:461–475.

90. Torrey, J., and A. Gray. 1841. “A” Flora of North America: Containing Abridged Descriptions of All the Known Indigenous and Naturalized Plants Growing North of Mexico; Arranged According to the Natural System. Wiley and Putnam.

91. Tucker, C. M., M. W. Cadotte, S. B. Carvalho, T. J. Davies, S. Ferrier, S. A. Fritz, R. Grenyer, M. R. Helmus, L. S. Jin, and A. O. Mooers. 2017. A guide to phylogenetic metrics for conservation, community ecology and macroecology. Biological Reviews 92:698–715.

92. Vilà, M., J. L. Espinar, M. Hejda, P. E. Hulme, V. Jarošík, J. L. Maron, J. Pergl, U. Schaffner, Y. Sun, and P. Pyšek. 2011. Ecological impacts of invasive alien plants: a meta- analysis of their effects on species, communities and ecosystems. Ecology letters 14:702–708.

93. Wang, G., Zhang, X., Yannelli, F., Li, J. J., Shi, S., Zhang, T., … & Jiang, L. (2024). The impact of species phylogenetic relatedness on invasion varies distinctly along resource versus non-resource environmental gradients. Journal of Applied Ecology, 61(4), 869–883.

94. Webb, C. O. 2000. Exploring the phylogenetic structure of ecological communities: an example for rain forest trees. The American Naturalist 156:145–155.

95. Webb, C. O., D. D. Ackerly, and S. W. Kembel. 2008. Phylocom: software for the analysis of phylogenetic community structure and trait evolution. Bioinformatics 24.

96. Webb, C. O., Ackerly, D. D., McPeek, M. A., & Donoghue, M. J. (2002). Phylogenies and community ecology. Annual review of ecology and systematics, 475-505.

97. Wiens, J. J., D. D. Ackerly, A. P. Allen, B. L. Anacker, L. B. Buckley, H. v Cornell, E. I. Damschen, T. Jonathan Davies, J. Grytnes, and S. P. Harrison. 2010. Niche conservatism as an emerging principle in ecology and conservation biology. Ecology letters 13:1310–1324.

98. Winter, M., Schweiger, O., Klotz, S., Nentwig, W., Andriopoulos, P., Arianoutsou, M., … & Kühn, I. (2009). Plant extinctions and introductions lead to phylogenetic and taxonomic homogenization of the European flora. Proceedings of the National Academy of Sciences, 106(51), 21721–21725.

99. Xu, J., Dang, H., Tian, T., Liu, S., Chai, Y., Liu, X., … & Chang, J. (2020). Human disturbance rather than habitat factors drives plant community assembly and diversity patterns in a semiarid region. Land Degradation & Development, 31(14), 1803–1811.

100. Zanne, A. E., D. C. Tank, W. K. Cornwell, J. M. Eastman, S. A. Smith, R. G. FitzJohn, D. J. McGlinn, B. C. O’Meara, A. T. Moles, and P. B. Reich. 2014. Three keys to the radiation of angiosperms into freezing environments. Nature 506:89–92.

