## Supplementary material for "Darwin’s Naturalization Conundrum explained by gradients of environmental stress and disturbance": Table S1

**Table S1.** Results from the global linear mixed-effects model for the unweighted mean distance of the non-native species relative to all native species (MDNS), the distance of the non-native species to its nearest native species (DNNS), and the distance to the single most abundant native species (DMANS) as response of elevation, plot type (interior or roadside) and their interaction. For elevation, we summarized the linear and quadratic terms of elevation using orthogonal polynomials. Significant p-values ($p < 0.05$) are indicated in bold.

| | MDNS | | | DNNS | | | DMANS | | |
|---|---|---|---|---|---|---|---|---|---|
| | $df_{num, den}$ | F-value | P-value | $df_{num, den}$ | F-value | P-value | $df_{num, den}$ | F-value | P-value |
| Vegetation Cover | 1, 981 | 23.610 | **<0.01** | 1, 918 | 4.367 | **<0.01** | 1, 981 | 21.413 | **<0.01** |
| Elevation | 2, 714 | 43.837 | **<0.01** | 2, 719 | 23.632 | **<0.01** | 2, 707 | 35.454 | **<0.01** |
| Disturbance | 1, 536 | 269.417 | **<0.01** | 1, 541 | 216.167 | **0.04** | 1, 518 | 168.782 | **<0.01** |
| Elevation: Disturbance | 2, 428 | 16.045 | **<0.01** | 2, 433 | 10.463 | **<0.01** | 2,409 | 12.143 | **<0.01** |

**Table S2.** Model comparison. We compared the global and regional linear and quadratic models based on the Akaike information criterion (AICc) and selected the model with the lowest AICc. The results from the Chi-square test (Pr (>Chisq)) highlighted in bold indicate that the AIC of the quadratic model was significantly lower than the AIC of the linear model.

| Model | AIC quadratic | AIC linear | Pr (>Chisq) |
|---|---|---|---|
| Global | 2120 | 2123 | **0.03** |
| Central Argentina | 346 | 346 | 0.11 |
| Southern Argentina | 270 | 274 | **0.02** |
| Australia (NSW) | 373 | 370 | 0.53 |
| Australia (Victoria) | 359 | 357 | 0.92 |
| Austria | 91 | 99 | **<0.01** |
| Central Chile | 138 | 139 | 0.08 |
| Southern Chile | 334 | 336 | 0.38 |

| | | | |
|---|---|---|---|
| China | 183 | 185 | **0.05** |
| Ecuador | 131 | 153 | **<0.01** |
| India | 145 | 160 | **<0.01** |
| Norway | 148 | 150 | **0.04** |
| Spain | 220 | 222 | **0.04** |
| Switzerland | 156 | 155 | 0.28 |
| USA (Hawaii) | 128 | 128 | 0.22 |
| USA (Montana) | 266 | 268 | 0.06 |
| USA (Oregon) | 363 | 379 | **<0.01** |

**Table S3.** The global linear mixed-effects model for weighted mean distance from non-native to native species (WMDNS) as a function of elevation, disturbance type (disturbed or undisturbed vegetation plots) and their interaction, as well as vegetation cover included as covariate. For elevation, we summarized the linear and quadratic term of elevation using orthogonal polynomials. Significant p-values ($p < 0.05$) are indicated in bold.

| | WMDNS | | |
|---|---|---|---|
| | $df_{num, den}$ | F-value | P-value |
| Vegetation Cover | 1, 965 | 25.91 | **<0.01** |
| Elevation | 2, 695 | 37.64 | **<0.01** |
| Disturbance | 1, 469 | 233.12 | **<0.01** |
| Elevation: Disturbance | 2, 360 | 21.06 | **<0.01** |

**Table S4.** Regional linear mixed-effects models for Weighted Mean Distance from the non-native to all Native Species (WMDNS) as a function of elevation, disturbance type (disturbed or undisturbed plots) and their interaction. For regions where a quadratic term of elevation was maintained ($df_{num}$= 2), we summarized the linear and quadratic term of elevation using orthogonal polynomials. Significant p-values ($p < 0.05$) are indicated in bold.

| **Region** | **Vegetation Cover** | | | **Elevation** | | | **Disturbance** | | | **Elevation: Disturbance** | | |
|---|---|---|---|---|---|---|---|---|---|---|---|---|
| | **df** | **F-value** | **P-value** | **df** | **F-value** | **P-value** | **df** | **F-value** | **P-value** | **df** | **F-value** | **P-value** |
| Central Argentina | 1, 102 | 13.24 | **<0.01** | 1, 54 | 43.93 | **<0.01** | 1, 50 | 21.59 | **<0.01** | 1, 51 | 9.76 | **<0.01** |
| Southern Argentina | 1, 70 | 1.21 | 0.28 | 2, 70 | 21.76 | **<0.01** | 1, 70 | 57.27 | **<0.01** | 2, 70 | 6.52 | **<0.01** |
| Australia (NSW) | 1, 94 | <0.01 | 0.98 | 1, 59 | 12.43 | **<0.01** | 1, 43 | 20.26 | **<0.01** | 1, 46 | 0.32 | **0.57** |
| Australia (Victoria) | 1, 93 | 17.42 | **<0.01** | 1, 93 | 53.82 | **<0.01** | - | - | - | - | - | - |
| Austria | - | - | - | 2, 24 | 12.15 | **<0.01** | 1, 24 | 0.54 | **0.46** | 2, 24 | 1.05 | **0.36** |
| Central Chile | 1, 29 | 0.89 | 0.35 | 1, 23 | 33.78 | **<0.01** | 1, 15 | 0.28 | **0.6** | 1, 15 | 0.02 | **0.87** |
| Southern Chile | 1, 90 | 5.64 | **0.02** | 1, 54 | 27.27 | **<0.01** | 1, 49 | 7.74 | **0.01** | 1, 55 | 0.3 | **0.58** |
| China | 1, 47 | 2.74 | 0.1 | 1, 47 | 0.38 | 0.54 | 1, 47 | <0.01 | **0.97** | 1, 47 | 0.38 | **0.54** |
| Ecuador | - | - | - | 2, 17 | 15.84 | **<0.01** | 1, 16 | 46.78 | <0.01 | 2, 16 | 2.77 | 0.09 |
| India | 1, 48 | 0.18 | 0.68 | 2, 22 | 50.34 | **<0.01** | 1, 30 | 0.35 | **0.56** | 2, 22. | 1.35 | **0.28** |
| Norway | 1, 34 | 3.43 | 0.07 | 2, 34 | 8.55 | **<0.01** | - | - | - | - | - | - |
| Spain | 1, 38 | 0.23 | 0.64 | 2, 41 | 2.45 | 0.09 | 1, 27 | 46.05 | **<0.01** | 2, 26 | 0.04 | **0.96** |
| Switzerland | 1, 33 | 0.44 | 0.51 | 1, 33 | 2.7 | 0.11 | 1, 33 | 0.35 | 0.55 | 1, 33 | 0.17 | 0.68 |
| USA (Hawaii) | 1, 35 | 22.83 | **<0.01** | 1, 36 | 8.63 | **<0.01** | - | - | - | - | - | - |
| USA (Montana) | - | - | - | 1, 83 | 113.87 | **<0.01** | 1, 83 | 9.92 | **<0.01** | 1, 83 | 3.41 | **0.06** |
| USA (Oregon) | 1,99 | 0.958 | 0.33 | 2, 99 | 123.51 | **<0.01** | 1, 99 | 34.75 | **<0.01** | 2, 99 | 6.75 | **<0.01** |

**Table S5**. Results from the global linear mixed-effects model for standardized Weighted Mean Distance for the native community (ses.MPD) as response of Elevation, disturbance type (disturbed or undisturbed plots) and their interaction. Significant p-values (p < 0.05) are indicated in bold.

| | Ses.MPD | | |
|---|---|---|---|
| | $df_{num, den}$ | F-value | P-value |
| Elevation | 2, 785 | 2.766 | 0.07 |
| Disturbance | 1, 465 | 29.646 | **<0.01** |
| Elevation: Disturbance | 2, 457 | 2.391 | 0.09 |

**Table S6.** Results from the regional linear mixed-effects models for standardized Mean Pairwise Distance (sMPD) as response of elevation, disturbance type (disturbed or undisturbed plots) and their interaction. For some of the regions, we summarized the linear and quadratic term of elevation using orthogonal polynomials. Significant p-values (p < 0.05) are indicated in bold.

| Region | Elevation | | | Plot type | | | Elevation: Plot type | | |
|---|---|---|---|---|---|---|---|---|---|
| | df | F-value | P-value | df | F-value | P-value | df | F-value | P-value |
| Central Argentina | 1, 59 | 0.418 | 0.52 | 1, 56 | 0.996 | 0.32 | 1, 58 | 0.635 | 0.43 |
| Southern Argentina | 2, 60 | 0.396 | 0.67 | 1, 53 | 1.329 | 0.25 | 2, 58 | 0.634 | 0.53 |
| Australia (NSW) | 1, 50 | 14.071 | **<0.01** | 1, 50 | 0.322 | 0.57 | 1, 49 | 1.428 | 0.24 |
| Australia (Victoria) | 2, 93 | 6.348 | **<0.01** | - | - | - | - | - | - |
| Austria | 1, 25 | 2.566 | 0.12 | 1, 8 | 1.398 | 0.27 | 1, 8 | 0.038 | 0.85 |
| Central Chile | 1, 19 | 1.168 | 0.29 | 1, 17 | 2.153 | 0.16 | 1, 18 | 4.961 | **0.04** |
| Southern Chile | 1, 60 | 0.055 | 0.82 | 1, 54 | 3.377 | 0.07 | 1, 56 | 0.157 | 0.69 |
| China | 1, 48 | 27.799 | **<0.01** | 1, 48 | 2.492 | 0.12 | 1, 48 | 0.777 | 0.38 |
| Ecuador | 2, 13 | 3.314 | 0.07 | 1, 13 | 1.366 | 0.26 | 2, 13 | 2.119 | 0.16 |
| India | 1, 26 | 1.101 | 0.3 | 1, 26 | 0.676 | 0.42 | 1, 26 | 2.071 | 0.16 |
| Norway | 2, 35 | 5.047 | **0.01** | 1, 38 | 9.384 | **<0.01** | 1, 37 | 1.037 | 0.31 |
| Spain | 1, 40 | 17.264 | **<0.01** | 1, 28 | 0.038 | 0.85 | 1, 30 | 4.234 | 0.05 |
| Switzerland | 1, 44 | 2.372 | 0.13 | 1, 24 | 0.061 | 0.81 | 1, 26 | 0.217 | 0.64 |
| USA (Hawaii) | 2, 28 | 15.32 | **<0.01** | - | - | - | - | - | - |
| USA (Montana) | 1, 64 | 4.073 | **0.05** | 1, 46 | 5.323 | **0.03** | 1, 49 | 6.685 | **0.01** |
| USA (Oregon) | 1, 52 | 0.342 | 0.56 | 1, 49 | 0.542 | 0.47 | 1, 50 | 0.865 | 0.35 |

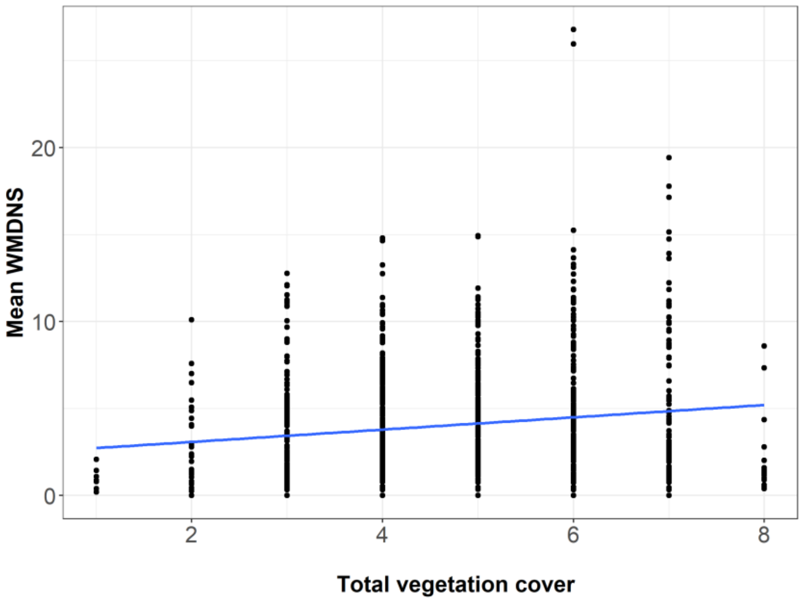


**Figure S1.** Global patterns of variation with total vegetation cover for Weighted Mean Phylogenetic Distance from the non-native to all Native Species (WMDNS).

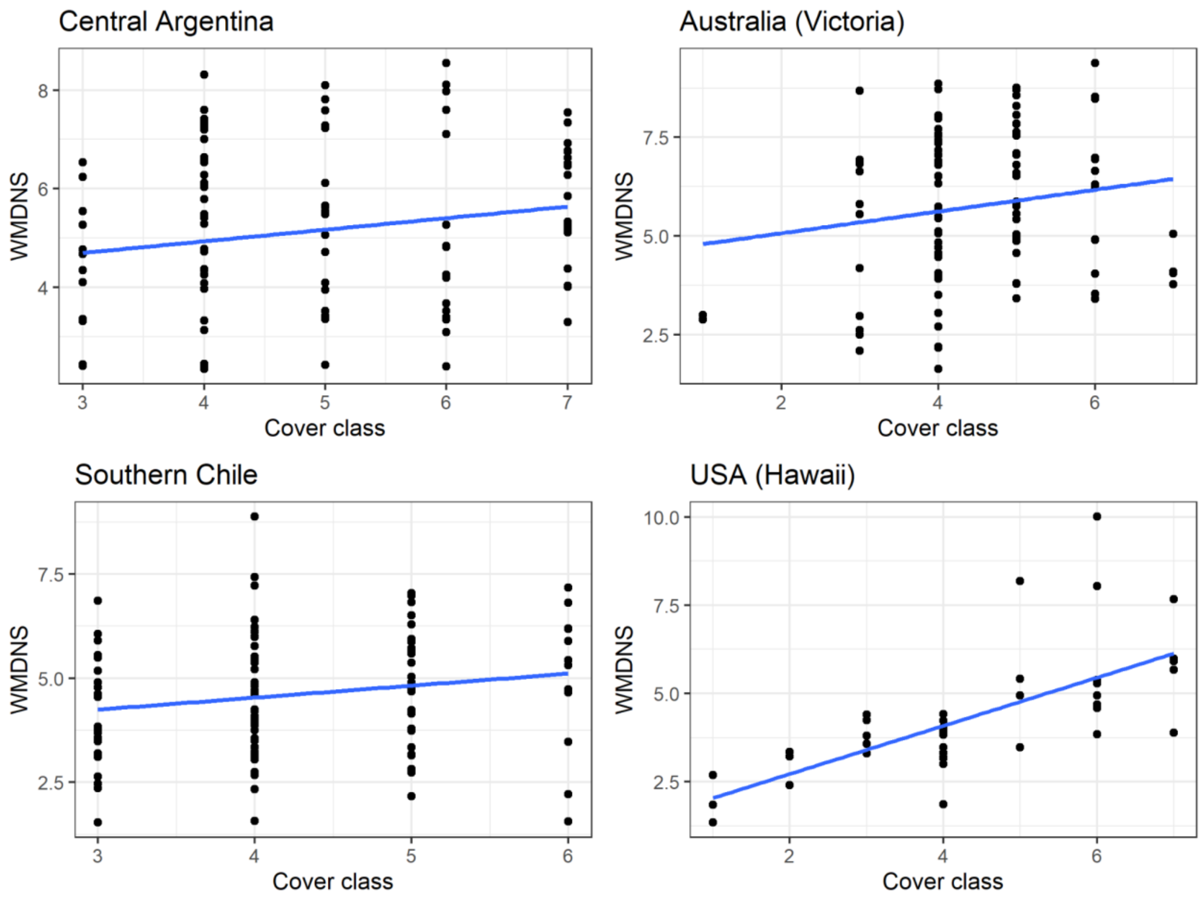

**Figure S2.** Regional patterns of variation with total vegetation cover for Weighted Mean Phylogenetic Distance from the non-native to all Native Species (WMDNS) for the regions where cover significantly affected phylogenetic distance.

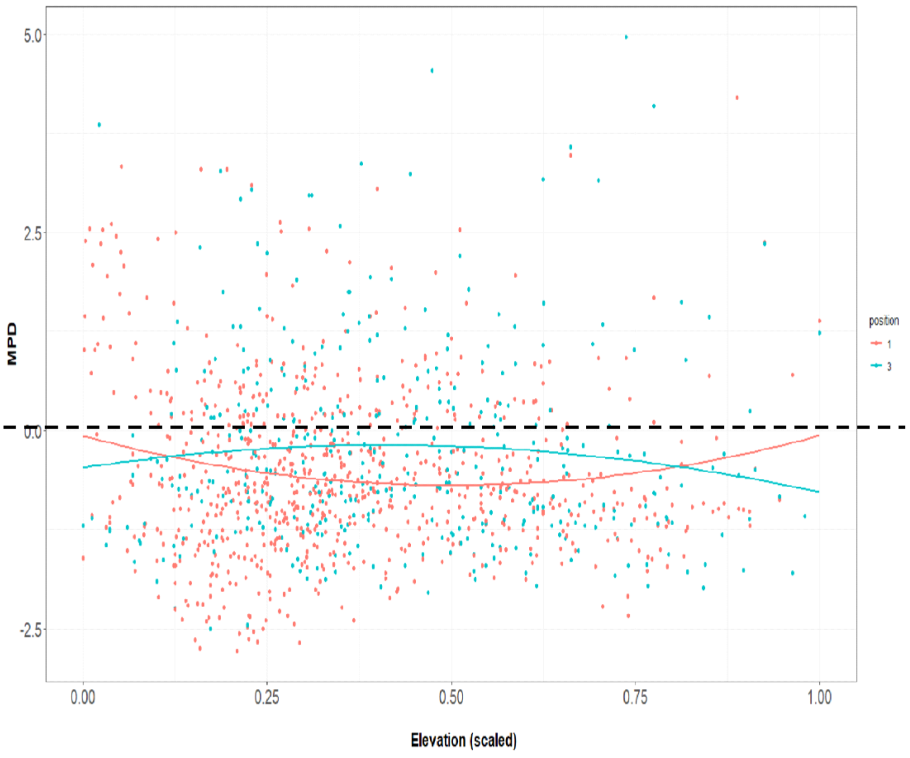


**Figure S3** Global pattern of elevational variation for standardized Mean Pairwise Distance (sMPD) of the native community in roadside plots (red dots and line) and interior plots (blue dots and line). Negative values indicate that the community is more clustered than expected by chance, while positive values indicate that the community is more overdispersed than expected.

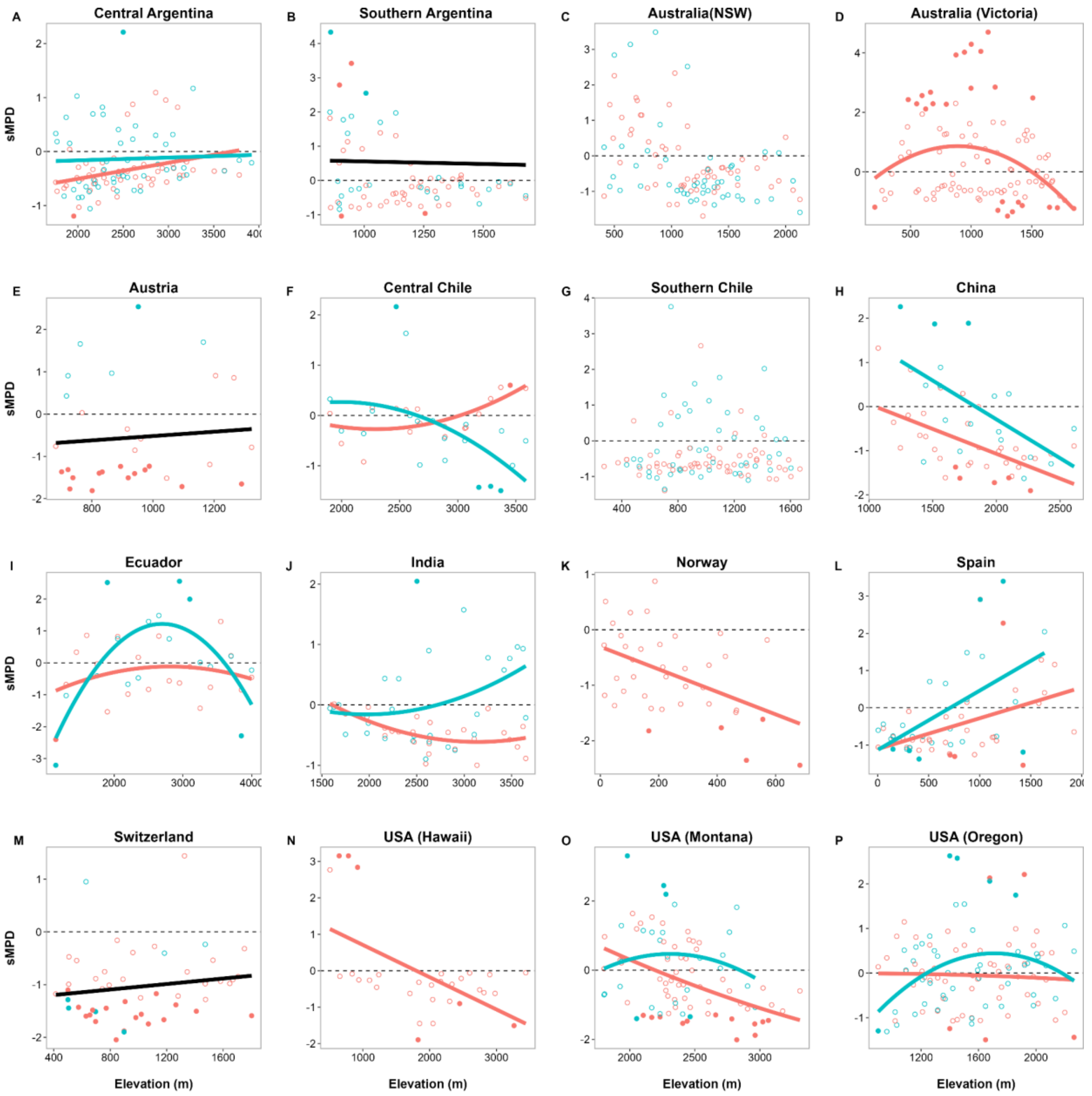


**Figure S4** Regional patterns of elevational variation for standardized Mean Pairwise Distance (sMPD) of the native community in disturbed plots ( red dots and lines) and undisturbed plots (blue dots and lines), black lines indicate the effect of elevation was the same for disturbed and undisturbed plots. Negative values indicate that the community is more clustered than expected by chance, while positive values indicate that the community is more overdispersed than expected by chance. Unfilled dots indicate that observed values were not significantly different from the predicted values.
